# The *Chlamydia* type III effector TarP alters the dynamics and organization of host cell focal adhesions

**DOI:** 10.1101/250563

**Authors:** António T. Pedrosa, Korinn N. Murphy, Ana T. Nogueira, Amanda J. Brinkworth, Tristan R. Thwaites, Jesse Aaron, Teng-Leong Chew, Rey A. Carabeo

**Author notes:** Corresponding author: Rey A. Carabeo, Ph.D., Department of Pathology and Microbiology, University of Nebraska Medical Center, Omaha, NE 68198-5900. Department of Pharmacology, University of North Carolina at Chapel Hill, Chapel Hill, NC.

## Abstract

The human pathogen *Chlamydia trachomatis* targets epithelial cells lining the genital mucosa. We observed that infection of various cell types, including fibroblasts and epithelial cells resulted in the formation of unusually stable focal adhesions that resisted disassembly induced by the myosin II inhibitor, blebbistatin. Super-resolution microscopy revealed in infected cells the vertical displacement of paxillin and FAK from the signaling layer of focal adhesions; while vinculin remained in its normal position within the force transduction layer. The candidate type III effector TarP which localized to focal adhesions during infection and when expressed ectopically, was sufficient to mimic both the reorganization and blebbistatin-resistant phenotypes. These effects of TarP, including its localization to focal adhesions required interaction with the host protein vinculin through a specific domain at the C-terminus of TarP. The consequence of *Chlamydia*-stabilized focal adhesions was restricted cell motility and enhanced attachment to the extracellular matrix. Thus, via a novel mechanism, *Chlamydia* inserts TarP within focal adhesions to alter their organization and dynamics.

## Introduction

Bacterial infection of mucosal epithelial cells triggers the antimicrobial defense strategy of cell exfoliation and apoptosis induction (reviewed in: Kim *et al*., 2010). The controlled extrusion of damaged host cells and colonizing pathogens requires the degradation of cell adhesion factors. In epithelial cells, focal adhesions and hemidesmosomes are primarily responsible for attachment to the extracellular matrix, and their assembly and turnover are exquisitely regulated at multiple levels, by kinases, phosphatases, protein-protein interactions, internalization of components, and degradation (Borradori and Sonnenberg, 1999; Geiger *et al*., 2001; Rosenblatt *et al*., 2001; Zaidel-Bar *et al*., 2007). Disruption of one or more of these regulatory processes alters the adhesion dynamics and properties of the cells.

One strategy employed by bacteria to neutralize exfoliation relies on the precise targeting of one or more components of the focal adhesion proteome. The best-characterized example is that of *Shigella*, which neutralizes epithelial extrusion to colonize the epithelium efficiently (Kim *et al*., 2009). It does so by delivering the OspE effector by the type III secretion system (T3SS). This protein reinforces host cell adherence to the basement membrane by interacting with integrin-linked kinase (ILK), a serine/threonine kinase that is part of the focal adhesome (Kim *et al*., 2009; Zaidel-Bar *et al*., 2007). A consequence of the OspE-ILK interaction is an increased surface expression of β1-integrin, which in turn promotes focal adhesion (FA) assembly. In addition, the OspE-ILK complex stabilizes the focal adhesions (FAs) by reducing phosphorylation of focal adhesion kinase (FAK) at a functionally important Tyr397 residue and of paxillin. Inhibition of both phosphorylation events has been shown to induce FA disassembly (Kim *et al*., 2009). Interestingly, some EPEC and EHEC strains, as well as *Citrobacter rodentium* possess the effector EspO, which shares strong homology with OspE (reviewed in Vossenkämper, Macdonald and Marchès, 2011; Morita-Ishihara *et al*., 2013). As such, it is conceivable that these pathogens also reinforce adherence of the infected epithelial cells to secure an infectious foothold. The EspZ effector of EPEC and EHEC has been shown to reduce cell death and detachment *in vitro* (Shames *et al*., 2010). EspZ binds the transmembrane glycoprotein CD98 and enhances its effect on β1-integrin signalling and cell survival via activation of FAK (Shames *et al*., 2010).It is possible that EspO and EspZ may cooperate to confer enhanced adhesion of the host epithelial cells to the extracellular matrix. Finally, through interaction with human carcino-embryonic antigen-related cell adhesion molecules (CEACAM), bacterial pathogens such as *Neisseria gonorrhoeae*, *Neisseria meningitidis*, *Moraxella catarrhalis*, and *Haemophilus influenzae* can activate β1-integrin signalling and inhibit epithelial cell detachment (reviewed in: Kim *et al*., 2010). Despite numerous examples of pathogens manipulating host cell adhesion, the details of these mechanisms remain uncharacterized.

Chlamydiae are obligate intracellular pathogens that are distinguished by their biphasic developmental cycle that alters between the infectious elementary body (EB), and the replicative, but non-infectious reticulate body (RB). At late time points, the non-infectious RBs convert back to EBs to produce infectious particles for the next round of infection. The entire intracellular growth cycle of *Chlamydia* takes ∼48–96 h and occurs within a membrane-bound inclusion, and most of it is spent in the non-infectious RB form. Thus, it is essential that the adhesion of the infected cells to the epithelium is sustained during chlamydial development to enable the differentiation of the non-infectious RBs to the infectious and stable elementary bodies (EBs) (reviewed in AbdelRahman and Belland, 2005). This means that *Chlamydia* must evade a host of anti-microbial defenses, including epithelial extrusion.

Previous works by Kumar and Valdivia (2008) and Heymann *et al*., (2013) described the loss of motility of *Chlamydia*-infected epithelial cells. Heymann *et al*., (2013) attributed this to the chlamydial inhibition of Golgi polarization that occurs at >24 h post-infection, leading to loss of directional migration. In this report, we offer an alternate and possibly complementary mechanism of FA stabilization, which could lead to an increase of host-cell adhesion to the extracellular matrix (ECM), thus culminating in previously reported loss of motility (Kumar and Valdivia, 2008; Heymann *et al*., 2013). Using quantitative confocal and live-cell imaging and super-resolution microscopy, we describe the various *Chlamydia* infection-dependent changes that occur to FAs, such as increased numbers, enhanced stability, and altered organization. We provide evidence implicating the T3SS effector TarP, and its interaction with the focal adhesion protein vinculin. We show that vinculin and its binding motif in TarP is required for the localization of the effector to focal adhesions, and their resistance to blebbistatin-induced disassembly. TarP localization to focal adhesions is also required for the displacement of the focal adhesion kinase and paxillin from their normal position within the integrin signaling layer. We also show that TarP alone was sufficient to restrict cell motility. Overall, the results indicate that *Chlamydia* has a dedicated mechanism of modulating focal adhesion dynamics, which may be linked to the maintenance of *Chlamydia* infection in a high-turnover tissue site.

## RESULTS

### Chlamydia infection enhances FA numbers

Cos7 cells were infected and 24 h post-infection (hpi), cells were fixed and prepared for indirect immunofluorescence imaging of paxillin-positive FAs. As shown in Figure 1, cells infected with *C. trachomatis* serovar L2, serovar D, serovar B, *C. caviae* GPIC, and *C. muridarum* (MoPn) consistently had greater numbers of FAs than mock-infected cells. We further explored the apparent infection-dependent increase in FA numbers using serovar L2, and observed enhanced FA numbers at 8 hpi that increased by 24 hpi. Next, we asked if the process is pathogen-directed. Specifically, we investigated if this effect on FAs required de novo protein synthesis by *Chlamydia*. Cos7 cells were infected with CtrL2 for the specified duration followed by treatment by the bacterial translation inhibitor, chloramphenicol (Cm). We observed that while an 8-h protein synthesis inhibition was not sufficient to prevent the effects on FA number, the 24-h Cm treatment reduced FA numbers of infected cells to the level of mock-infected control (Figure 2). The results indicate that the latter phase of focal adhesion alterations requires either the *de novo* synthesis of new proteins by *Chlamydia* or replenishment of effectors packaged in the metabolically quiescent EBs. These effectors are delivered by the type III secretion system early in infection, prior to differentiation to the vegetative reticulate body form, when they gain the ability for macromolecular synthesis.

**Figure 1.**
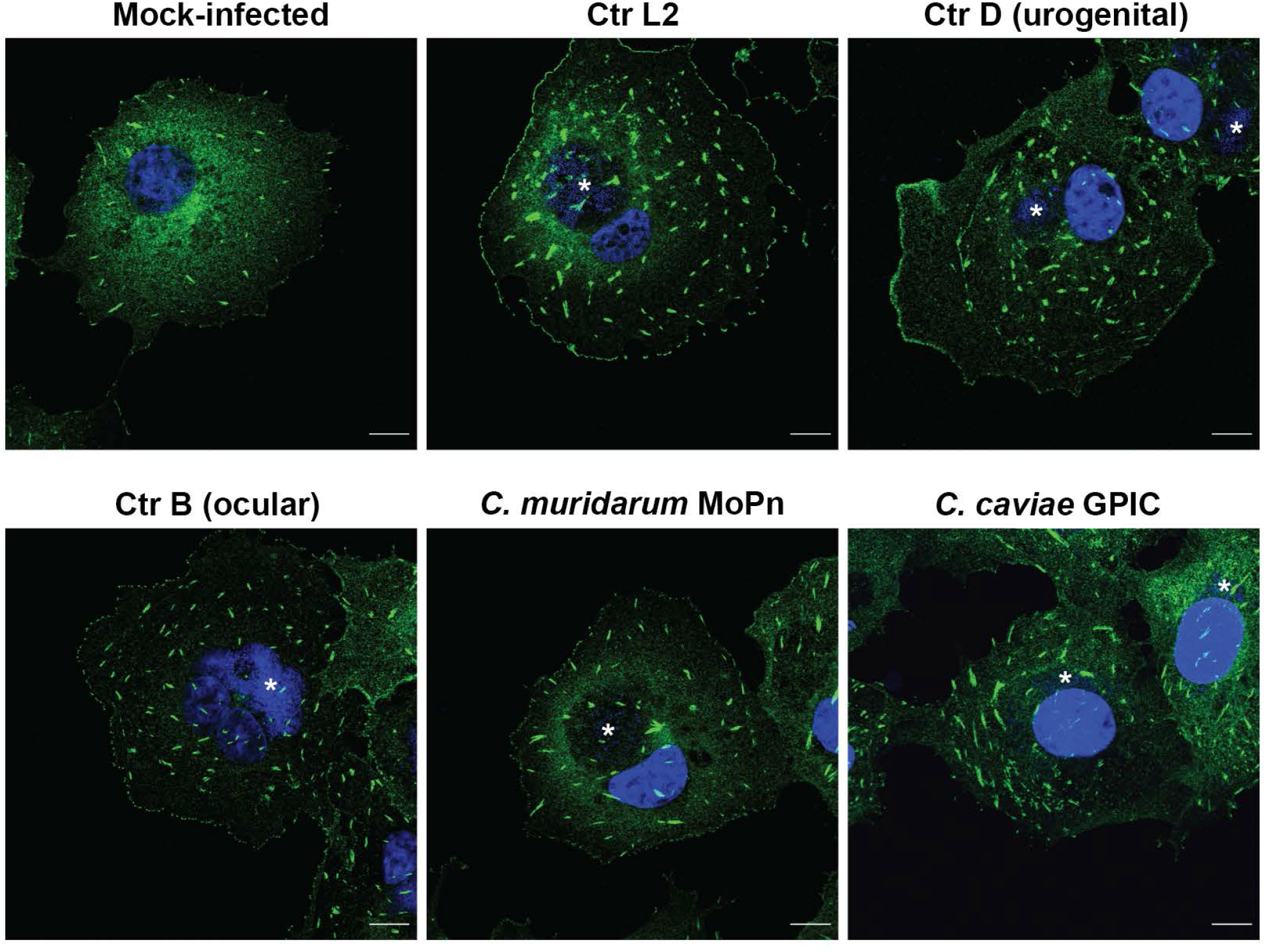
*C. trachomatis*-infected Cos7 cells exhibit increased focal adhesion numbers. (A) Cos7 cells infected with the indicated chlamydial strain/species or mock-infected were monitored at 24 hpi, and focal adhesions visualized by immunostaining for paxillin.

**Figure 2.**
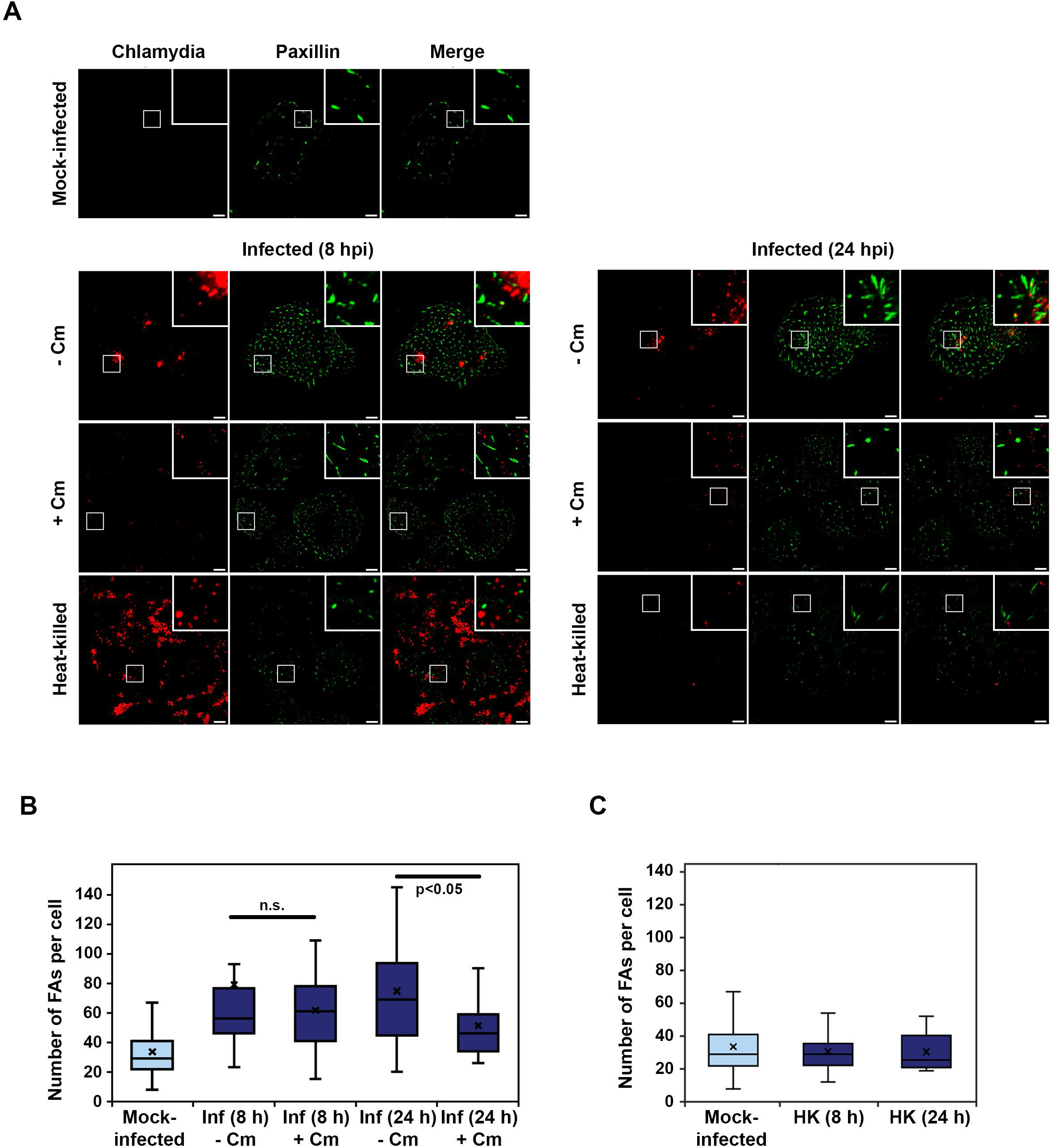
Infection-dependent increase in focal adhesion numbers requires *de novo* chlamydial protein synthesis. (A) Cos7 cells infected with *C. trachomatis* serovar L2 were mock- or chloramphenicol-treated at the start of infection for either 8 or 20 h. Focal adhesions were visualized by immunostaining for paxillin (green). Scale bar length 10µm. (B) Focal adhesions were counted using the particle tracker plug-in in NIH ImageJ. Analysis revealed a lack of effect of the 8-h Cm treatment on FA numbers, while the 24-h treatment yielded a statistically significant decreasein FA numbers. Data are represented as box-and-whisker plots. Whiskers represent the lowest and highest data point still within 1.5 times the interquartile range. The light blue asterisks indicate significant difference relative to the mock-infected control (Wilcoxon rank sum test * = p<0.05). The black cross shows the average for each experimental sample.

A marked difference was the increased numbers of FAs at the interior relative to the cell periphery. Focal adhesion maturation is associated with movement away from the cell periphery and towards the center (Smilenov, et al. 1999). FAs at the interior of the cell either mature to become stable fibrillar adhesions to promote cell attachment or disassemble during migration (Smilenov, et al. 1999; Dumbauld *et al*., 2010; reviewed in: Nagano *et al*., 2012). We speculated that the increased numbers of interior FAs arose from infection-dependent stabilization. To assess stability, we took advantage of the enhanced turnover of FAs in the presence of the myosin II-specific inhibitor, blebbistatin (Straight *et al*., 2003). FA stability is dependent on tension within and between focal adhesions (Chrzanowska-Wodnicka and Burridge, 1996; Pasapera *et al*., 2010; Carisey *et al*., 2013). This tension is largely provided by the contractile action of the molecular motor myosin II on stress fibers (SF); and its inhibition by blebbistatin consistently leads to FA disassembly and altered motility (Wang *et al*., 2008; Dumbauld *et al*., 2010, Liu *et al*., 2010). We evaluated the relative resistance of FAs to a 60-min treatment with 10 μM blebbistatin in the context of infection with CtrL2 at 20 hpi. As shown in Figure 3 of representative samples, SFs in the mock-infected cell disassembled, with the simultaneous disappearance of FAs marked with paxillin. In contrast, FAs in the CtrL2-infected cells remained. Infection did not prevent blebbistatin-induced disassembly of stress fibers. Taken together, the results point to a dedicated mechanism in *Chlamydia trachomatis* and perhaps other chlamydial species to stabilize focal adhesions. In addition, focal adhesion stabilization in infected cells did not require stress fibers.

**Figure 3.**
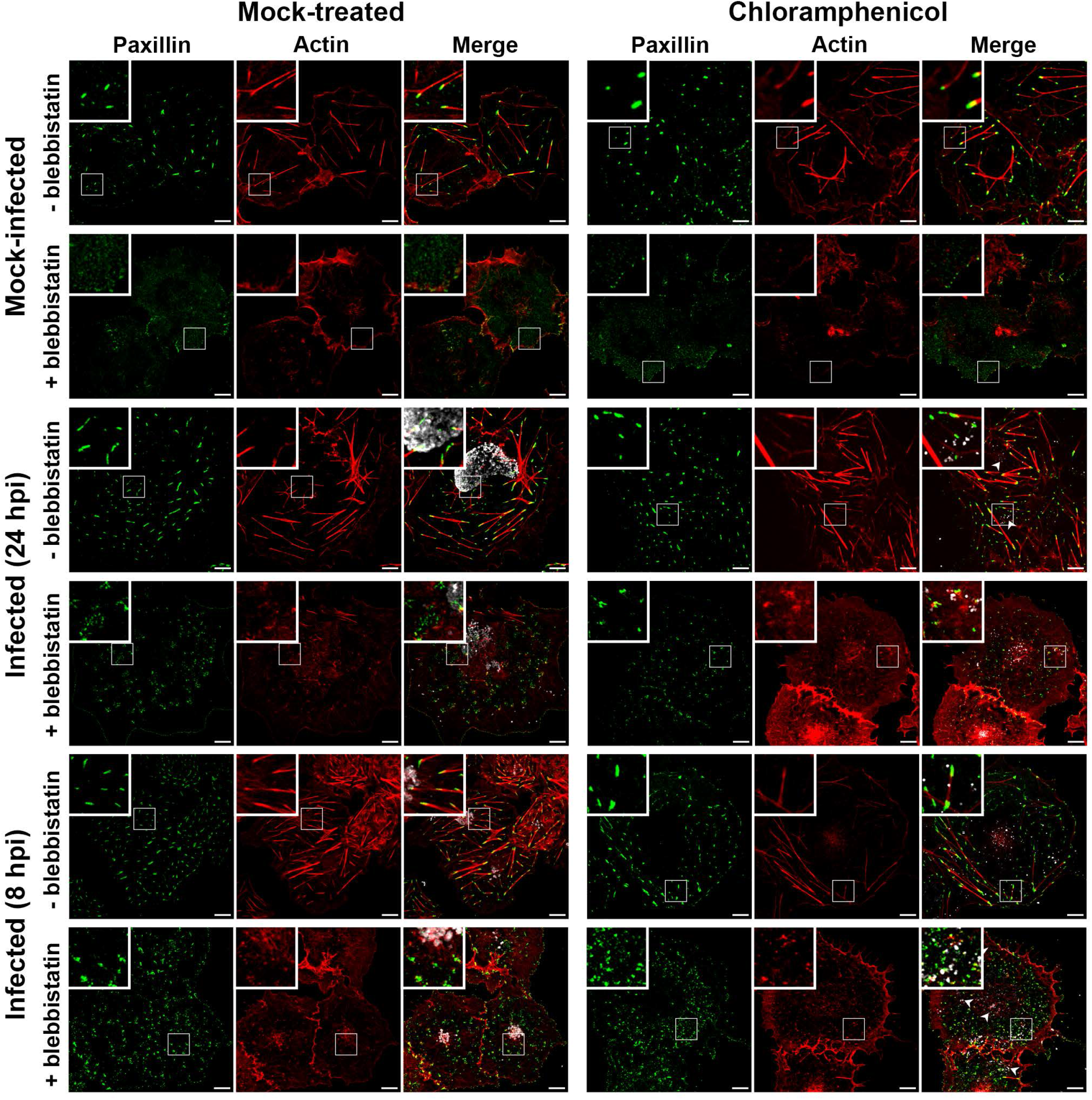
Focal adhesions of *Chlamydia* infected cells are resistant to blebbistatin. Cos7 cells were mock-infected or infected with CtrL2 for 24 h or 8 h. Cells were fixed and stained for the focal adhesion marker paxillin (green), F-actin (red) and human convalescent serum for *C. trachomatis* (white). Cells were also mock- or pre-treated with chloramphenicol (Cm) followed by infection of live EBs. Cm treatment was maintained for the duration of the experiment. Blebbistatin 10 μM was introduced during the last hour of infection. Cells without blebbistatin treatment showed clear F-actin stress fibres and paxillin-labeled focal adhesions. While both structures were lost in mock-infected cells, infected cells retained the focal adhesions. This characteristic was sensitive to Cm inhibition of *de novo* chlamydial protein synthesis. Scale bar = 10µm.

### The type III effector TarP localizes to focal adhesions in a vinculin-dependent manner

The chlamydial type III effector TarP has been implicated in the invasion process during infection of non-phagocytic cells. Specifically, TarP translocation by the elementary bodies contributes to the actin remodeling that is required for uptake of the pathogen (Clifton *et al*., 2005; Jewett *et al*., 2006; Lane *et al*., 2008; Thwaites *et al*., 2014). Interestingly, TarP was also reported to have a role in increased resistance of infected cells to apoptosis, raising the possibility that this effector has post-invasion function (Mehlitz *et al*., 2010). Consistent with this idea is the continuous presence of the protein throughout infection (Clifton *et al*., 2004). In addition, there was sustained presence of TarP translocated to the cytosol as indicated by its reactivity to the anti-phosphotyrosine 4G10 antibody under conditions that prevented further synthesis of this protein, i.e. Cm treatment (Figure S1). During invasion, TarP localizes to sites of chlamydial adhesion at the plasma membrane (Clifton *et al*., 2004). If TarP has a role post-invasion, we expect TarP to be found at sites where it exerts its function. Immunostaining with a rabbit polyclonal antibody to C. *trachomatis* serovar L2 (CtrL2) TarP of infected mouse embryonic fibroblasts (MEFs) revealed specific staining of focal adhesions (FA), in addition to punctae within inclusions, which are likely to be the bacteria (Figure 4). Uninfected cells consistently exhibited diffused background immunofluorescence signal, illustrating specificity of the antibody to TarP localized to FAs.

**Figure 4.**
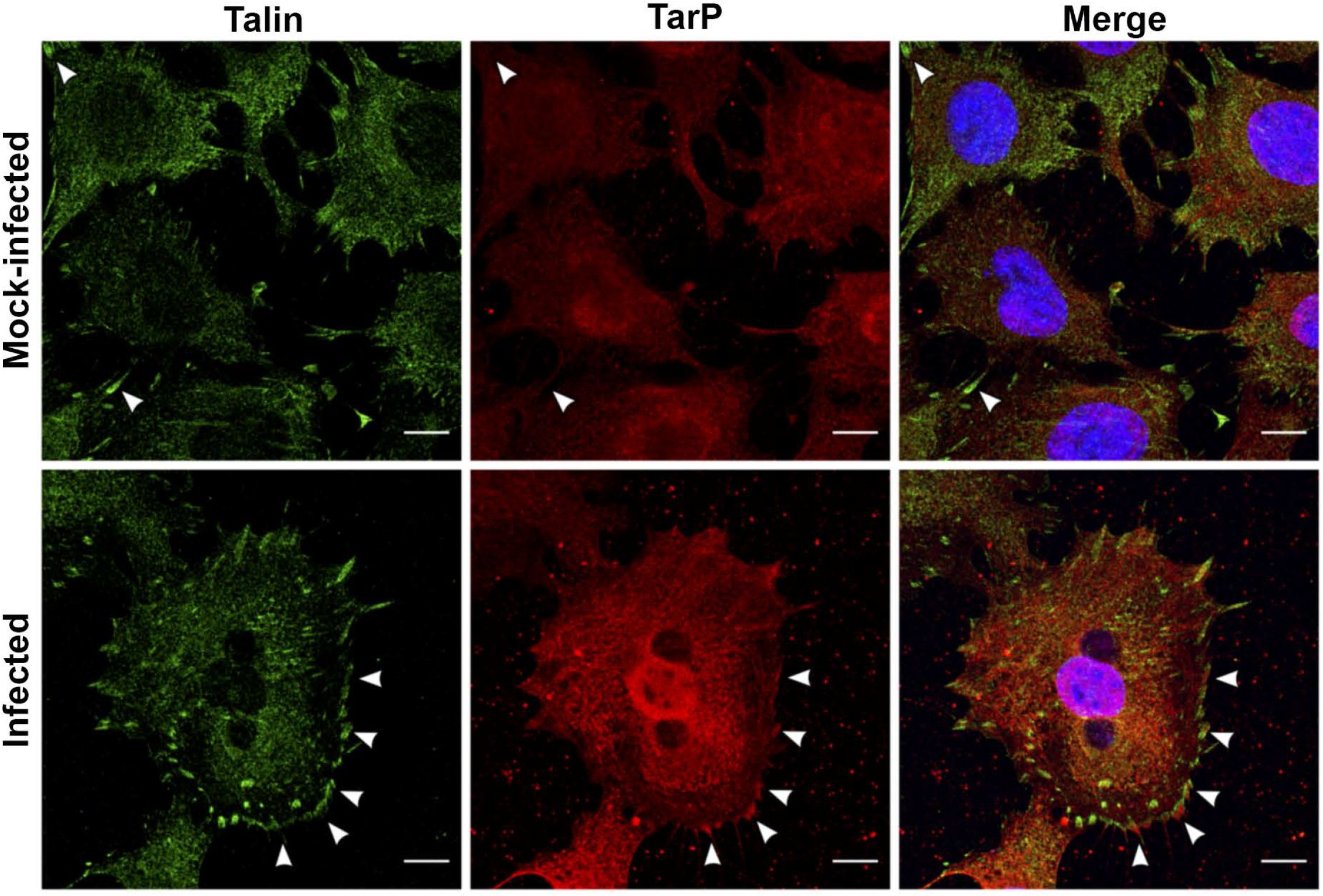
The type III effector TarP localizes to focal adhesions during *Chlamydia* infection. (A) CtrL2-infected MEF cells were immunostained for a rabbit polyclonal antibody to TarP and a mouse monoclonal antibody to talin. Inclusions were visualized by staining with DAPI. TarP localized to talin-positive FAs (arrowheads) in infected, but not in mock-infected controls. Scale bar = 10µm.

We then sought to determine if ectopically expressed TarP would yield a similar subcellular localization to FAs. TarP and its deletion derivatives shown in Figure 5A were fused to the fluorescent protein mTurquoise2, and ectopically expressed in MEFs. TarP has multiple domains that resemble motifs for protein-protein interaction and signaling, including a repeated 50-amino acid domains that is tyrosine-phosphorylated by Src family kinases and the Abl kinase, actin-binding domains, a leucine-aspartate (LD) domain recognized by the focal adhesion kinase (FAK), and vinculin-binding domains (VBD) (Clifton *et al*., 2005; Jewett *et al*., 2006, 2010; Lutter *et al*. 2010; Mehlitz *et al*. 2010; Jiwani *et al*., 2013; Thwaites *et al*., 2014, 2015; Braun *et al*., 2019). All have been demonstrated in invasion-related actin-recruitment assays to be functional(Lane *et al*., 2008; Thwaites *et al*., 2014, 2015). We created various C-terminal deletion constructs of TarP fused to mTurquoise for heterologous expression in MEFs, as well as a TarP derivative lacking the proline-rich domain (PRD) to minimize non-specific aggregation of the protein. Transfected cells were counterstained with a monoclonal antibody to paxillin to visualize FAs. As shown in Figure 5B, colocalization of TarP with the focal adhesion marker paxillin required the LD and VBD motifs. Note that paxillin localization was observed for both full-length TarP and TarP *Δ*PRD (Figure S2)

**Figure 5.**
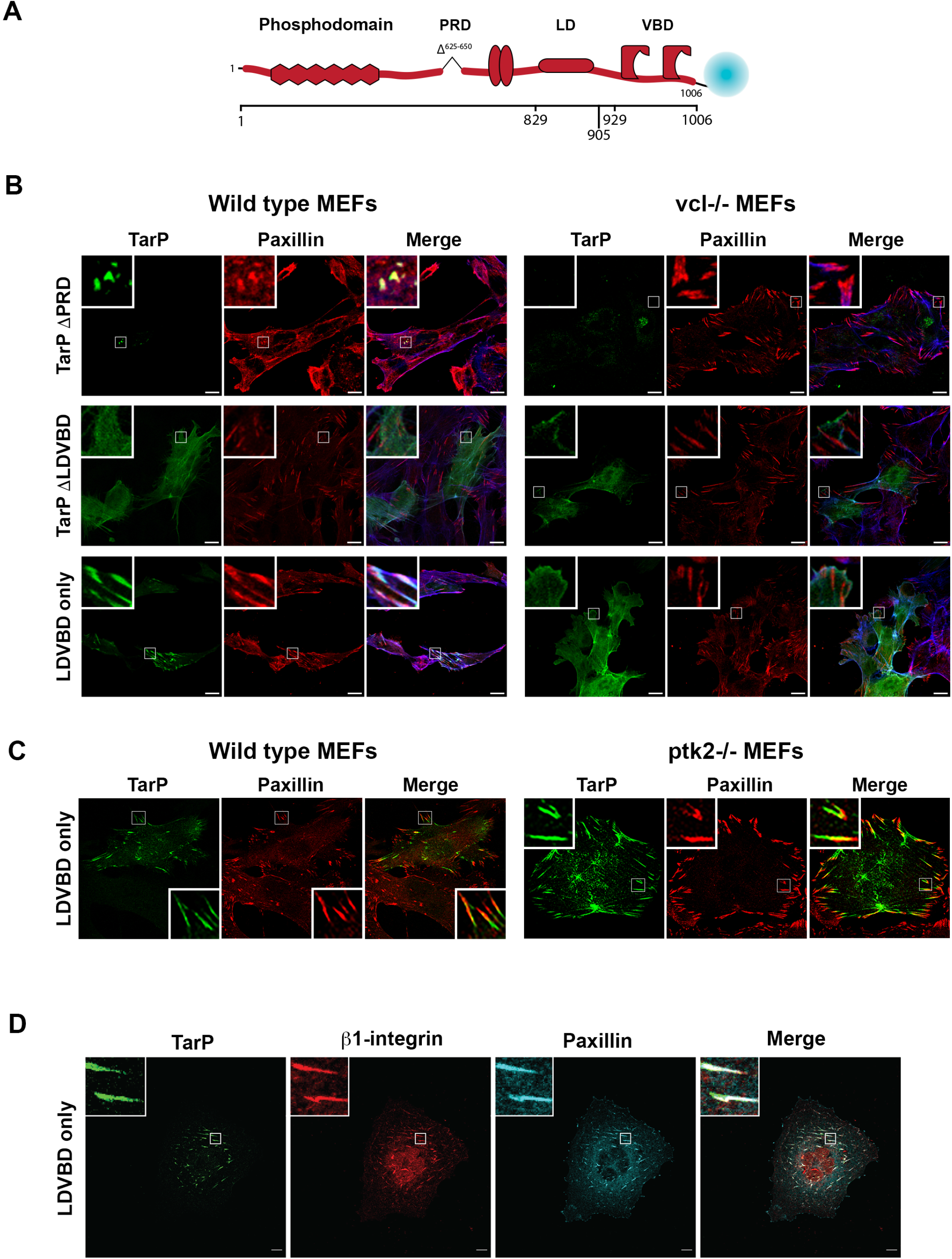
The focal adhesion localization of TarP requires its LDVBD domain and the host protein vinculin, but not FAK. (A) Representation of *C. trachomatis* effector protein TarP and its known domains fused to mTurquoise2 fluorescent protein. (B) Wild type or vinculin-knockout MEFs were transfected with different mTurquoise2-tagged TarP constructs (green) and imaged by confocal microscopy to evaluate colocalization with paxillin (red) at focal adhesions. Phalloidin was used to stain F-actin (blue). Cells were transfected for 20 h, at which time the cells were fixed and processed for immunofluorescence staining for paxillin. (C) In a parallel experiment, wild type and FAK1-deficient MEFs were transfected to express Flag-HA-LDVBD and stained for flag (green). Colocalization with paxillin (red) was assessed by confocal microscopy. (D) Ectopically expressed LDVBD localizes to β1-integrin and paxillin-positive focal adhesions. Scale bar = 10µm.

We previously reported that the LD and VBD domains were recognized by FAK and vinculin, respectively (Thwaites *et al*., 2014, 2015). They are distinct non-overlapping domains that interacted with their respective binding partners independently. Therefore, we evaluated if the loss of either of the binding partner (e.g. FAK or vinculin) would result in the loss of FA localization. To address the functional relevance of these interactions, albeit in a post-invasion context, the LDVBD construct was expressed in wild type, *vcl*−/− (vinculin), or *ptk2*−/− (FAK) MEF knockout mutants. We observed FA localization of mTurquoise-LDVBD in wild type MEFs, but not in *vcl*−/− MEFs (Figure 5B). The loss of FAK did not affect the focal adhesion localization of LDVBD (Figure 5C). Together the data indicated that TarP localization to FAs required the host protein vinculin, likely through its interaction with the VBD domain. FAK was dispensable in this regard. The TarP-positive subcellular structures were verified as FAs based on β1-integrin staining using a monoclonal antibody specific to the conformationally active form of the receptor (Figure 5D).

### The ectopic expression of TarP is sufficient to increase focal adhesion numbers and confer vinculin-dependent resistance to blebbistatin

We observed increased paxillin-positive FAs in TarP-transfected cells, provided that the TarP construct retained the VBD domain. Images from the FA localization experiments in Cos7 cells were re-analyzed by quantifying the number of focal adhesions per cell in cells transfected with the mTurquoise vector alone, full-length TarP, or LDVBD (Figure 6A). Paxillin-positive structures in transfected cells were counted in NIH ImageJ, and the data is illustrated as box-whisker plots in Figure 6B. We observed statistically significant increases in focal adhesion numbers in cells expressing TarP *Δ*PRD and LDVBD. These results suggested that the ability of TarP to localize to FAs, which was mediated by the LDVBD domain of TarP was linked to its effects on FA numbers.

**Figure 6.**
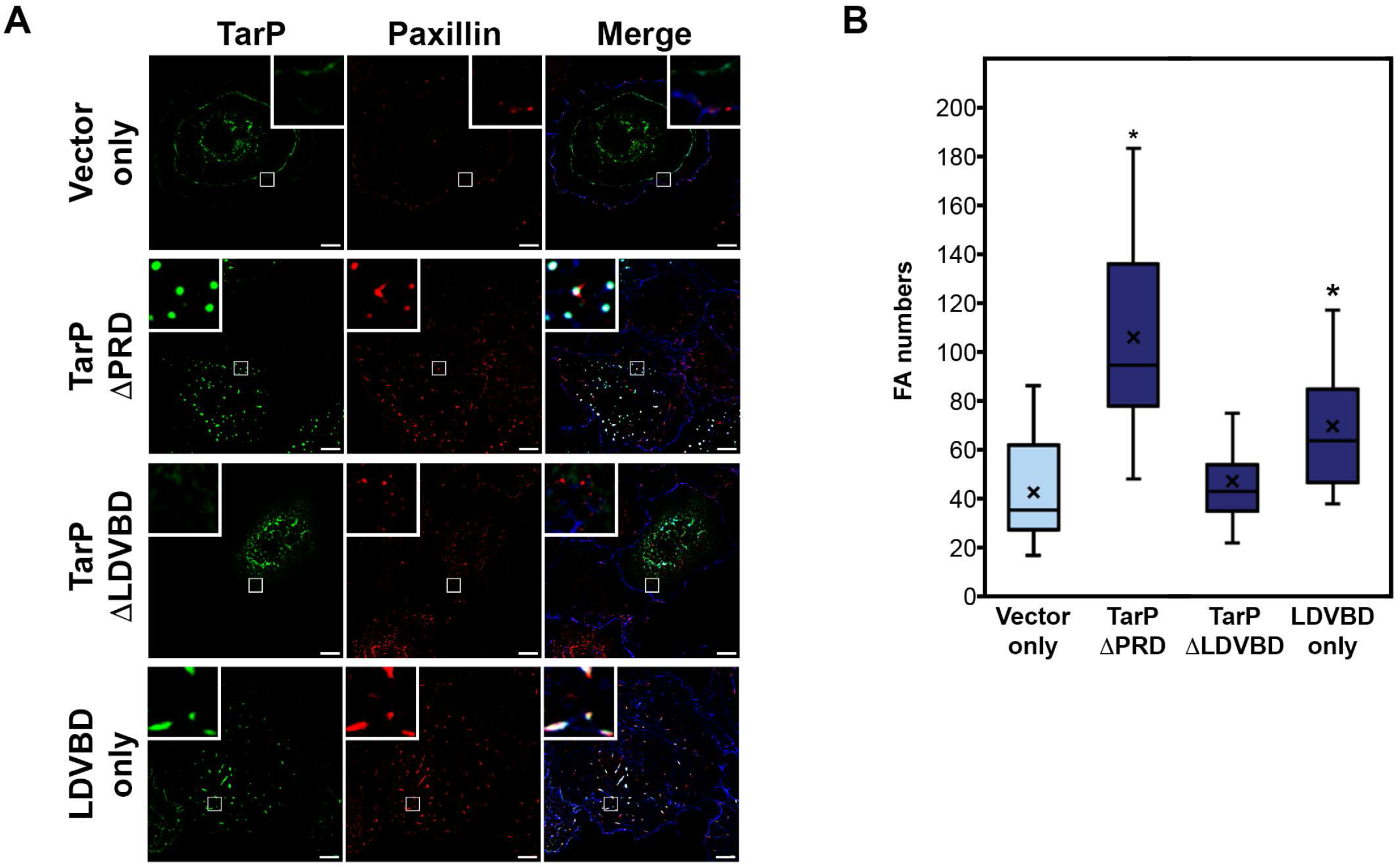
The ectopic expression of TarP LDVBD is sufficient to increase focal adhesion numbers. (A) Cos7 cells expressing different deletion derivatives of TarP or vector only were processed for immunofluorescence with anti-paxillin antibody to visualize focal adhesions. Representative images are shown. Scale bar = 10 µm. (B) Focal adhesion numbers were counted using the particle counting plug-in in NIH ImageJ. Data are focal adhesion number per cell and illustrated as box-whisker plot. Whiskers represent the lowest and highest data point still within 1.5 times the interquartile range. For statistical analyses the Wilcoxon Rank sum test was used to determine significance when compared to vector-only control (* = p<0.05).

We also evaluated the effect of ectopically expressed LDVBD on the stability of focal adhesions in fixed cells. As illustrated in Figure 7A, *Chlamydia*-infected wild type MEFs retained paxillin-marked FAs after 60 min of treatment with 10 uM blebbistatin, while mock-infected cells lost them. In contrast, infection of the *vcl*−/− MEFs failed to inhibit blebbistatin-induced disassembly of focal adhesions, highlighting the crucial role of the host protein vinculin in FA stability. Using these results as reference, we investigated if the LDVBD domain was sufficient to confer a similar level of resistance to blebbistatin-induced disassembly, and if it did so in a vinculin-dependent manner. LDVBD transfection of wild type MEFs led to the retention of FAs after the 60-min treatment with blebbistatin, while the *vcl*−/− MEFs lost these structures despite LDVBD expression (Figure 7B). Therefore, we concluded that vinculin plays an important role in TarP-dependent stabilization of FAs. However, we could not determine if the apparent stabilizing role of vinculin is due to the FA localization of TarP or an alteration of its activity as a result of its interaction with TarP.

**Figure 7.**
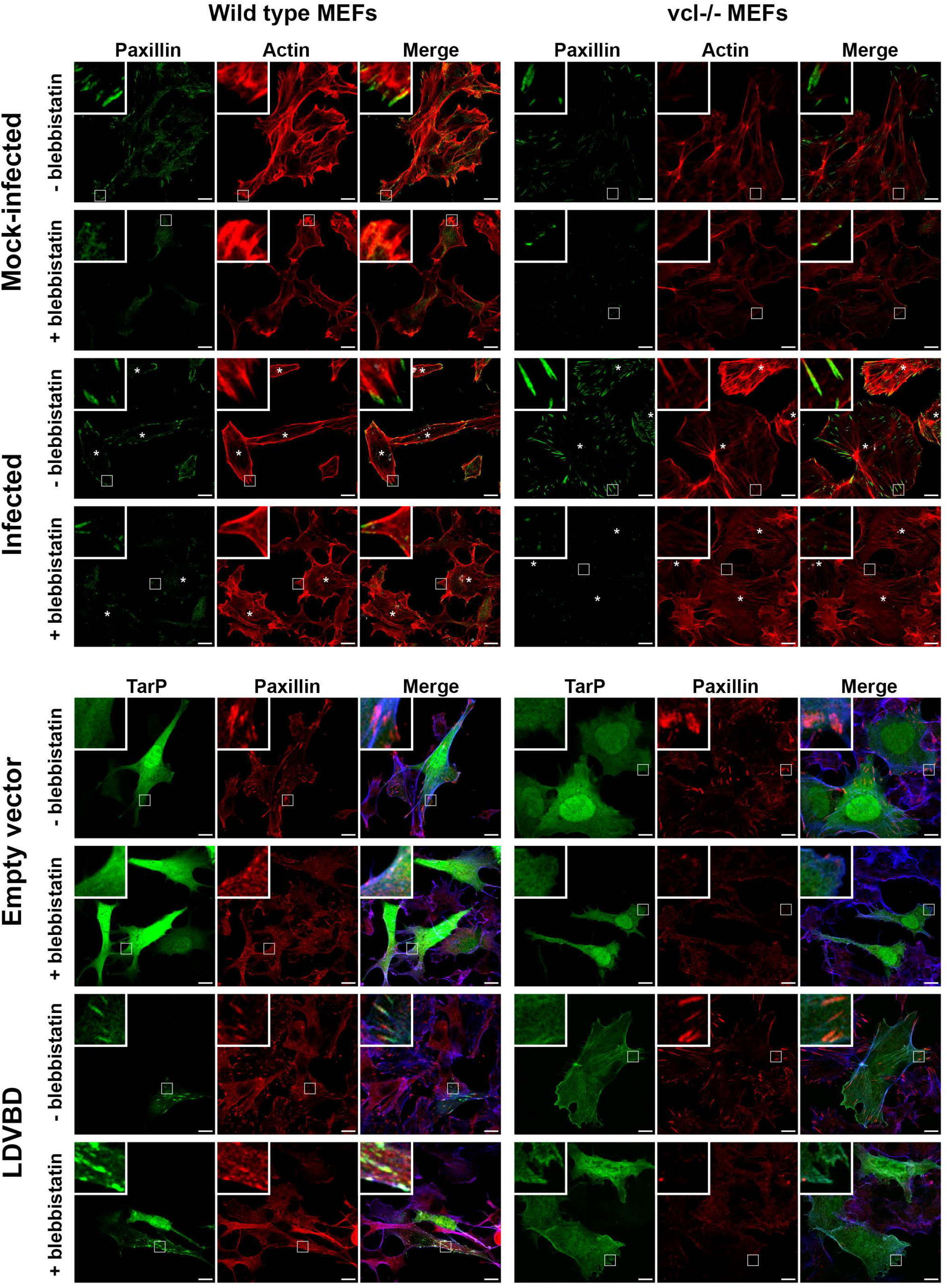
The LDVBD domain of TarP and the host protein vinculin are required for focal adhesion resistance to blebbistatin treatment. (A) Wild type or vinculin-knockout MEFs were infected with C. trachomatis serovar L2. Cells at 24 hpi were mock-treated or treated for 60 min with 10 μM of blebbistatin. The cells were then processed for immunofluorescence staining for paxillin (green) and actin (red). Retention or loss of focal adhesions were monitored. Focal adhesions were only resistant to blebbistatin-induced disassembly if the cell was infected and expressing vinculin. (B) In a parallel experiment, wild type or vinculin-knockout MEFs were transfected for 20 h with the empty vector or LDVBD-mTurquoise2 fusion protein. During the last hour, cells were either mock- or blebbistatin-treated. Cells were processed to visualize paxillin (red), LDVBD (green), and actin (blue; shown in composite images). LDVBD was sufficient to confer resistance to blebbistatin-induced disassembly to focal adhesions. Resistance also required vinculin. Scale bar = 10 µm.

### Infection disrupts the stratified organization of focal adhesions, a phenotype mimicked by the ectopic expression of TarP

Focal adhesions are organized into distinct strata termed the integrin layer, the signaling layer, which harbors paxillin and FAK amongst others, and the force transduction layer that contains vinculin, talin, and other mechanosensitive proteins. At the highest layer, actin and actin-associated proteins, such as *α*-actinin and myosin II are found (Betzig *et al*., 2006; Kanchanawong *et al*., 2010). Given the profound effect of infection and TarP ectopic expression on FA stability, we investigated using interferometric photoactivated localization microscopy (iPALM) their effects on FA organization. The localization of paxillin, FAK, and vinculin, all fused to mTurquoise2 were monitored in mock-infected, CtrL2-infected, TarP full-length, or LDVBD-transfected cells (Figure 8). Image analysis revealed dramatic reorganization of FAs with regards to paxillin and FAK. In control samples, paxillin, FAK, and vinculin were found 50.5, 40.4, and 71 nm from the bottom of the cell, respectively, consistent with previous findings (Kanchanawong *et al*., 2010). However, paxillin and FAK shifted upwards to 176.9 and 96 nm, respectively in infected cells, while no change in location was observed for vinculin (Figure 8A). The state of infection of cells analyzed are shown in Figure S3.

**Figure 8.**
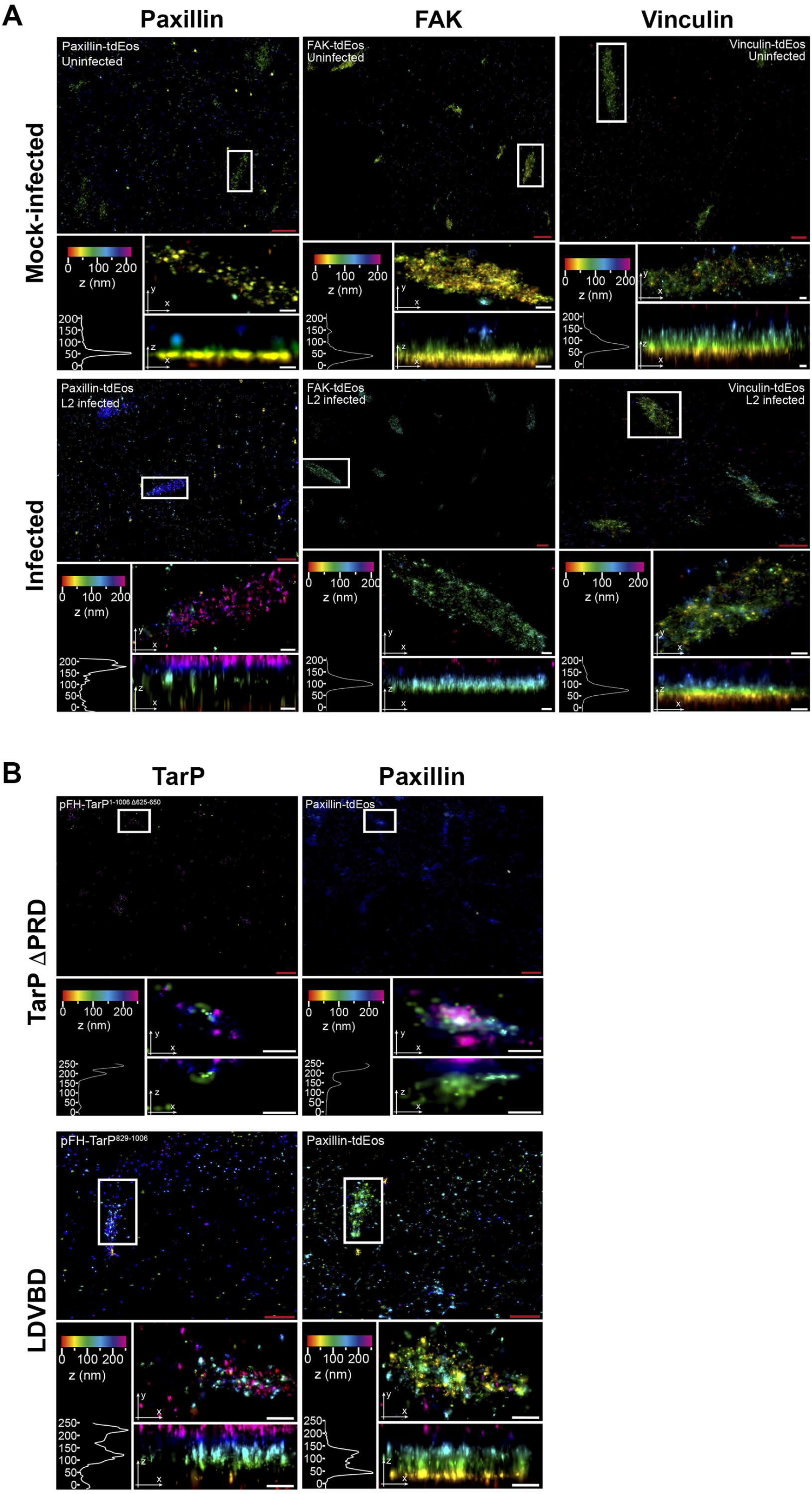
TarP-targeted focal adhesions display altered nanoscale architecture. (A) Cos7 cells pre-transfected with paxillin-tdEos, FAK-TdEos, or Vinculin-TdEos, and grownon gold fiducial coverslips were mock- or C. trachomatis-infected for 24 h. The cells were fixed and processed for iPALM imaging. Representative images are shown. For each sample, multiple panels are provided. The top panel shows the top view of area around the focal adhesion of interest (white border). The middle panel displays a top view of the focal adhesion indicated by the white border. The bottom panel shows the side view and corresponding z histograms. Note the significant shifts in paxillin and FAK localization, but not vinculin. (B) Cos7 cells were co-transfected with paxillin-tdEos and either TarP *Δ*PRD or LDVBD only by electroporation. The cells were seeded on gold fiducial coverslips, and processed for iPALM at 20 h post-transfection. Description of each panel is as above in (A). Note the significant shift in the location of paxillin within the TarP-positive focal adhesions. The various colors indicate the distance (z-coordinates) from the gold fiducial marker, (e.g. z = 0 nm; red). Red scale bar = 1µm. White scale bar = 200 nm.

Expression of TarP *Δ*PRD also caused a shift (240.2 nm) in paxillin localization (Figure 8B). LDVBD expression caused a noticeable shift (47 nm, with a second peak at 130 nm), but to a lesser degree than full-length TarP. Interestingly, vinculin was not affected by either infection or TarP *Δ*PRD (or LDVBD) expression, which points to the specific disruption of FA organization by *Chlamydia*.

### Cell motility is restricted in Chlamydia-infected or TarP-expressing cells

It was previously reported that *Chlamydia*-infected cells were restricted in their motility, and this was attributed to the inability of infected cells to establish front-rear polarity due to Golgi fragmentation induced by the pathogen. We decided to reinvestigate the loss of motility of infected cells by focusing on focal adhesion dynamics. The decision of the cell to migrate or adhere involves the regulation of focal adhesion stability in response to external cues, such as chemoattractants and extracellular matrix (ECM) stiffness. First, we verified that *Chlamydia*-infected mouse embryo fibroblasts were severely limited in their ability to migrate, relative to mock-infected control cells (Movie S1). The manual tracking plugin of ImageJ was utilized to obtain cell trajectory tracks for the motility assay. Cells were tracked using the position of the nucleus over time. The coordinate data was input into ibidi’s chemotaxis and migration tool to obtain velocity and distance measurements. Velocity measurements revealed a 1.5-fold decrease in the mean rate of migration of infected cells (Figure 9A **and** 9C).

**Figure 9.**
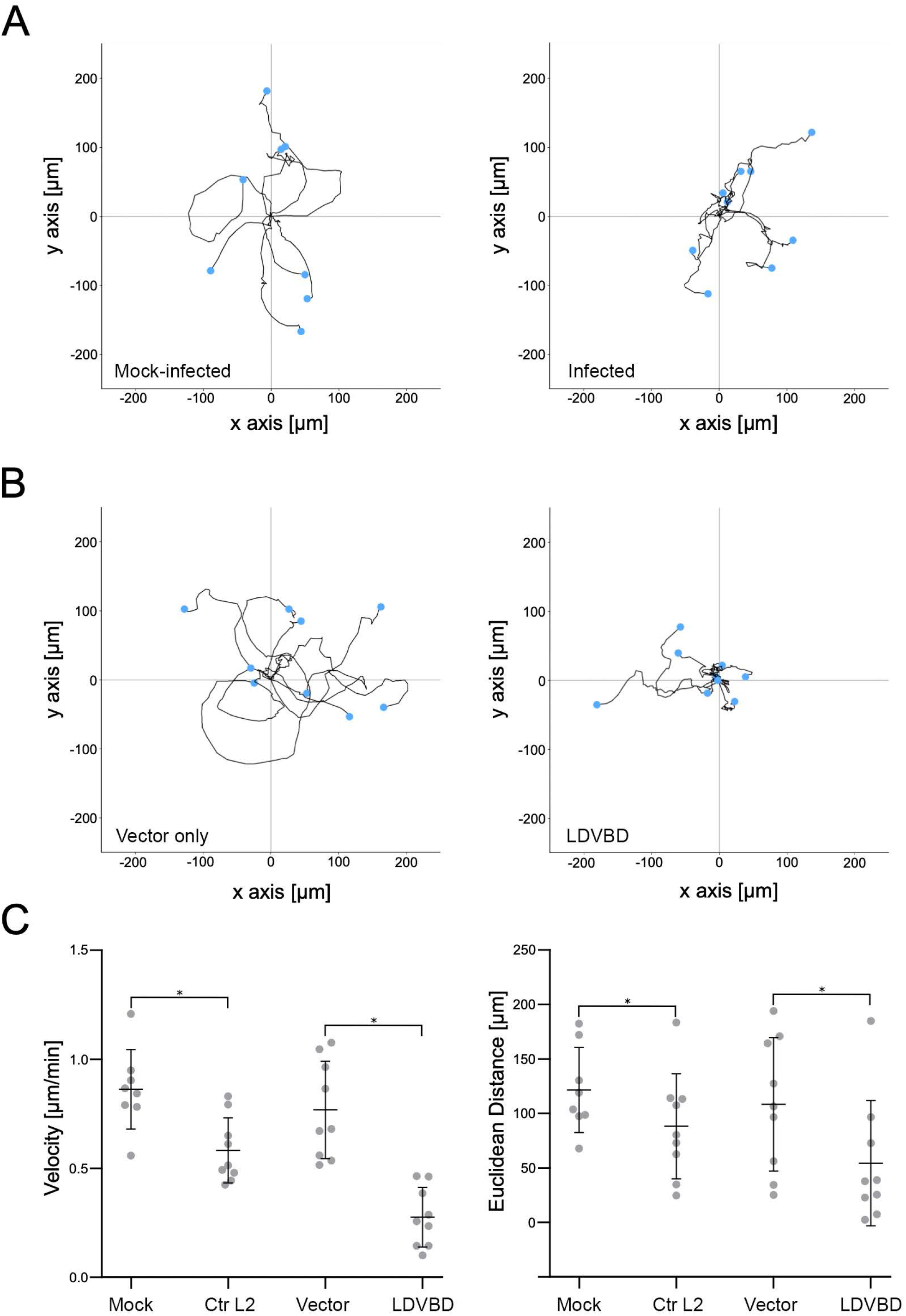
The LDVBD domain of TarP is sufficient to inhibit cell migration. MEFs that were mock-infected, Chlamydia-infected, vector-only-transfected, or LDVBD-transfected were seeded within ibidi µ-slide live cell imaging chambers. Time-lapse imaging was performed every 10 minutes for 10 h to evaluate cell motility. (A) For analysis of the infection experiments, a 5-h imaging window common to both mock- and *Chlamydia*-infected samples was chosen that maximized the number of cells that remained within the field of view. Cells were tracked using the manual tracking function in NIH ImageJ, and the cell trajectory was traced and plotted with the starting points assigned to the origin. (B) Analysis of the transfection experiment was in a common 10-h imaging window. Data were acquired and plotted as in (A). (C) Velocity and Euclidean distance traveled were calculated for each cell from each experimental group. Values were plotted as dot-plots with mean ± S.D. indicated by the bars. Statistical significance was calculated using the Wilcoxon Rank Sum test. * indicates p < 0.05.

Mouse embryonic fibroblasts were transfected to ectopically express mTurquoise2 vector only or the LDVBD domain. Cell migration was monitored in the DIC channel with fluorescence images taken at the end of the motility assay (**Movie S1**; Figure S4). To quantify motility, the cells were tracked as described above, with accompanying velocity calculations. As shown in Figure 9A **and** 9B, transfected cells were significantly restricted in their motility relative to mock-transfected controls. Both distance and velocity of LDVBD-transfected cells were further restricted to those of infected cells (Figure 9C) indicating that inhibition of cell migration by *C. trachomatis* could be accounted for fully by TarP overexpression. The enhanced inhibition of migration distance and velocity in transfected cells may have been due to increased levels of LDVBD when compared to levels present during infection.

### Infection by Chlamydia trachomatis but not ectopic expression of TarP confers resistance to detachment by mild trypsinization

Exfoliation of epithelial cells from the infected epithelium has been reported in rodent models of ocular and genital infection; and both reports speculated the involvement of neutrophil-derived proteases in the process (Ramsey *et al*., 2005; Lacy *et al*., 2011). We evaluated the resistance of infected epithelial cells to detachment by 0.025% trypsin, and monitored for cell rounding by time-lapse imaging at 1-min intervals for 30 min (Movie 1). Uninfected HeLa cells started detaching by 7 min post-trypsinization, while *C. trachomatis* L2-infected cells remained attached through the duration of imaging (30 min C. *trachomatis* post-trypsinization). To evaluate if TarP ectopic expression would be sufficient to resist detachment, the cells were transfected for 24 h to overexpress (LDVBD-mTurquoise2). Monolayers were imaged under fluorescence microscopy at 0, 15, and 30-min after trypsinization. If TarP LDVBD overexpression was sufficient to induce detachment resistance, we would expect an enrichment in remaining adherent cells of those expressing LDVBD-mTurquoise2 than cells expressing mTurquoise2 only. Percentage values for both LDVBD-mTurquoise2 and vector-only samples were 11.9% vs. 11.0% prior to trypsinization. We obtained the following for LDVBD vs. vector-only; 10.2% vs. 10.3% (15 min) and 8.7% vs. 11.0% (30 min). No statistically significant differences were found between LDVBD and vector only for either trypsinization time point. These results indicated that, while TarP is able to reduce cell motility, it was not sufficient to confer resistance to detachment by mild trypsinization. We interpret this to mean that additional changes to focal adhesions, possibly mediated by additional chlamydial factors are required.

## Discussion

All chlamydial species, to varying extents exhibit tropism to epithelial cells, and thus likely evolved to counteract cell extrusion associated with the normal cycle of turnover of the epithelium, or as an anti-microbial mechanism to limit dissemination and eliminate infection. The latter may also involve polymorphonuclear (PMN) cells, which secrete proteases to degrade adhesion structures of epithelial cells. Thus, *Chlamydia*, with its biphasic developmental cycle that involves a temporary loss of infectivity, is subjected to a very strong selective pressure to acquire mechanisms to inhibit epithelial cell extrusion. A large portion of the developmental cycle is spent in the non-infectious form, and thus, it is crucial to the survival of the pathogen to inhibit extrusion of host epithelial cells before *Chlamydia* could convert to the infectious form. Here, we demonstrated that adhesion of infected cells is enhanced via the action of TarP, an effector protein conserved in the genus *Chlamydia*, and the host cell protein vinculin.

Our data collectively point to FAs as targets for modulation by *Chlamydia*. The type III effector TarP plays a role in this modulation. The mechanism involves the vinculin-dependent localization of TarP to FAs resulting in increased stability as indicated by their resistance to disassembly by the myosin II inhibitor blebbistatin. We also made the novel observation of FA reorganization in *Chlamydia*-infected or TarP-expressing cells, with paxillin and FAK displaced from the signaling layer. A pressing question is whether the reorganization is the cause or the effect of enhanced stability of TarP-targeted FAs. Considering the reported relative stability of FAs in FAK-depleted cells, it is possible that the displacement of FAK by TarP could be analogous to a loss-of-function. A more comprehensive investigation of FA disorganization by TarP is required to define the exact mechanism of FA stabilization by this chlamydial effector. We did not observe any change to tyrosine autophosphorylation at position 397; and we cannot exclude the possibility that despite the apparent phosphorylation level, FAK displacement might negatively affect interactions with signaling proteins that recognize and bind this motif, like the Src kinase. The progression of tyrosine phosphorylation along the FAK protein by Src is crucial to the role of FAK in disassembling FAs. Therefore, restricted FAK-Src interaction as a consequence of FAK displacement in infected cells remains a possibility. Paxillin was similarly displaced in both infected and TarP-expressing cells, which likely disrupts protein-protein interactions and signaling related to paxillin within FAs. To our knowledge, FA reorganization to the extent that we observed in infected cells has not been reported, pointing to a possible novel mechanism of regulating FA dynamics.

While TarP, specifically the LDVBD domain was sufficient to drive FA stability to inhibit cell motility, it was unable to mimic the resistance of infected cells to detachment by mild trypsinization, which was meant to replicate polymorphonuclear cell (PMN)-mediated extrusion of infected epithelial cells. This would be consistent with the involvement of additional chlamydial factors that may mediate various aspects of focal adhesion characteristics, in addition to numbers. Additional factors may facilitate progression of maturation, possibly to fibrillar adhesions. Another might be the infection-dependent production of extracellular matrix components by the host cell, including collagen. Increased deposition of collagen underneath infected cells would likely influence FA stability by engaging integrins and inducing “outside-in” FA-stabilizing signals. Thus, *Chlamydia* might have multiple cooperating mechanisms ensuring the strong adhesion of its host cell, highlighting the importance of a mechanism to counteract epithelial cell extrusion.

A pressing question is the biological significance of FA stabilization by this pathogen. We speculate that this might be one mechanism by which *Chlamydia* neutralize extrusion of epithelial cells. It is known that this process could limit infection dissemination. Shedding of epithelial cells from uropathogenic *E. coli* (UPEC)-infected bladder is thought to reduce bacterial burden, facilitating resolution of infection (Mysorekar *et al*., 2002). The intestinal pathogen *Shigella* possesses an effector, OspE that modulates epithelial cell attachment to facilitate the pathogen’s cell-to-cell spread. Indeed, loss of OspE resulted in a significant decrease in virulence (Kim *et al*., 2009, 2010). Epithelial cells of the genital tract or the ocular mucosa also experience higher rates of turnover and are constantly replenished (Kuwabara, Perkins and Cogan, 1976; Crosson, Klyce and Beuerman, 1986; Anderson, Marathe and Pudney, 2014). Epithelial extrusion is a complex process that involves, not only promoting detachment of the cells from the ECM, but also modulating interactions with neighboring cells via disassembly of intercellular junctions (Rosenblatt, Raff and Cramer, 2001; Gudipaty and Rosenblatt, 2017). Another possibility is increased resistance to apoptosis. Cell detachment is associated with anoikis, a programmed cell death associated with loss of adherence (Frisch and Francis, 1994; Gudipaty *et al*., 2018). Focal adhesions provide anti-apoptotic signals (Frisch *et al*., 1996), and *Chlamydia* stabilization of these structures could promote apoptosis resistance.

An interesting question is the means by which epithelial cell extrusion is triggered. Is it part of the normal cell turnover during tissue remodeling/renewal, or is it linked to pathogen recognition? The shedding of bladder epithelial cells during UPEC infection requires the expression of bacterial Type 1 pili, which is a potent pathogen-associated molecular pattern (PAMP) that is recognized by the toll-like receptor 4 (TLR4) (Mysorekar *et al*., 2002), raising the intriguing possibility of a direct link between regulation of cell adhesion dynamics and pathogen recognition. Various chlamydial species are recognized by toll-like receptors expressed on epithelial cells e.g. TLR2, TLR3, TLR4, and TLR9 (Derbigny, Kerr and Johnson, 2005; O’Connell *et al*., 2006; Shaw *et al*., 2011; Derbigny *et al*., 2012; Pan *et al*., 2017; Carrasco *et al*., 2018; Kumar *et al*., 2019), which may be indicative of epithelial detachment being linked to pathogen recognition.

In addition to the demonstration that all tested *Chlamydia* species exhibited enhanced FA stability, the presence of a type III effector with FA modulating function highlights the importance of this process to *Chlamydia*; and yet, this aspect of Chlamydia-host cell interaction is relatively understudied. The availability of a *Chlamydia* mutant lacking TarP and an efficient method to image epithelial cell extrusion in the genital tract of animal models of infection are critical to advancing studies in this area.

## Materials and Methods

### Cell culture

Cos7 (ATCC CRL-1651), NIH3T3 (kindly supplied by Hector Aguilar-Carreño ATCC CRL-1658) and HeLa 229 (ATCC CCL-2.1) Mouse embryonic fibroblasts (MEFs) *vcl*^−/−^ and matched HeLa 229 MEFs *vcl*^+/+^ (Marg *et al*., 2010) (were kindly provided by Dr. Wolfgang Ziegler Hannover Medial School).cells were culture using Dulbecco’s Modified Eagle Medium (DMEM) (Thermofisher scientific, 11960-085). Media were supplemented with 10% fetal bovine serum (Sigma, F0804-500ML), 2mM L-glutamine, and 10μg/ml gentamicin. The human keratinocytes HaCaT cells (kindly supplied by Dr. Kristin M. Braun) were cultured in 3 parts DMEM and 1 part Ham’s F-12 Nutrient Mix (Thermofisher scientific 11765054), supplemented with 10% fetal bovine serum (Sigma, F0804-500ML), 2 mM L-glutamine, 10μg/ml gentamicin, insulin (Sigma I9278-5ML) and hydrocortisone cholera toxin EGF (HCE) cocktail. *Chlamydia trachomatis* serovar L2 (L2/434/Bu) was propagated in HeLa 229. EBs were harvested by discontinuous density gradient centrifugation in gastrografin (Bracco Diagnostics), as previously described (Thwaites *et al*., 2014).

### *Chlamydia* infections

Cells were infected with *Chlamydia trachomatis* serovar L2 (L2/434/Bu, CtrL2) at the multiplicity of infection MOI of 5, for 20 h, and of 25, for 8 h, in ice cold serum-free DMEM. Cells were centrifuged at 1000 rpm for 5 min at 4 °C to synchronize the infection. After centrifugation, the inoculum was replaced with warm DMEM supplemented with 10% fetal bovine serum, 2 mM L-glutamine, and 10μg/ml gentamicin. In parallel, a mock-infected control was made following the same protocol but without *Chlamydia* infectious particles. Infection by other strains/serovars was as follows. Cos7 cells were grown on glass coverslips. Cells were infected with *Chlamydia trachomatis* serovar L2, serovar D, serovar B, *Chlamydia muridarum* (MoPn), or *Chlamydia caviae* (GPIC) at an MOI of 5 for 24 hours. Cells were centrifuged at 500 x g for 15 minutes at 4°C to synchronize the infection. A mock-infected control was made following the same protocol but without *Chlamydia* infectious particles. Prior to infection with Serovar D, cells were pre-treated with 1x DEAE-Dextran for 15 minutes at room temperature. Pre-treatment was followed by two washes with 1x HBSS and replacement with DMEM to continue the infection.

### Immunostaining

Cells were grown on fibronectin coated coverslips (Neuvitro, GG-12-fibronectin) for the duration of the experiment. At the pre-determined, time cells were rinsed with Hank’s Balanced Salt Solution (HBSS) (Thermofisher scientific, 14025-100) and fixed using 4% paraformaldehyde (PFA) in PBS pH 7.4 (Gibco, 14190-094) for 20 min at room temperature. The fixed cells were then permeabilized using PBS with 0.2% Triton X-100. Subsequently, permeabilized cells were incubated with 1% BSA (Sigma, A9418) in PBS for 30 min at room temperature to block non-specific antigen binding. Cells were then incubated with the primary antibodies overnight at 4°C with rocking. The primary antibodies used in this study were rabbit polyclonal antibody against FAK phosphorylated at tyrosine 397 (pFAK-Y397) (Abcam, ab4803), rabbit monoclonal antibody paxillin (Abcam, ab32084), mouse monoclonal antibody vinculin (Abcam, ab18058), rat monoclonal 9EG7 against the active form of β1-integrin (BD Biosciences, 553715), mouse monoclonal Flag-tag antibody (Cell Signalling, 8146S) mouse monoclonal antibody *Chlamydia* LPS (Abcam, ab62708) and convalescent human sera. Afterwards cells were incubated with appropriate fluorescently conjugated secondary antibodies and, when specified, with DAPI (Roche, 10236276001) and Alexa flour 488 phalloidin stains, for 1 hr at room temperature, with rocking. In this study, the following secondary antibodies were used: goat anti-rabbit Alexa flour 488 (Thermofisher Scientific, A11008), goat anti-rabbit Alexa flour 633 (Thermofisher Scientific, A21071), goat anti-mouse Alexa flour 594 (Thermofisher Scientific, A11005), goat anti-human Alexa flour 647 (Thermofisher scientific A-21445). Following staining, the coverslips were mounted with Mowiol, and visualized in ZEISS LSM 710 confocal microscope, in the Microscopy and Histology Core Facility at the University of Aberdeen, or the Leica SP8 confocal microscope in Washington State University Integrative Physiology and Neuroscience advance image equipment. FIJI software (Schindelin *et al*., 2012; Schneider, Rasband and Eliceiri, 2012) was used to generate the final images.

To localize endogenous TarP to focal adhesions, Mouse embryonic fibroblasts (MEFs) grown on glass coverslips were infected with *Chlamydia trachomatis* L2 at an MOI of 10 for 20 hours. Cells were centrifuged at 500G for 15 minutes at 4°C to synchronize the infection. A mock-infected control was made following the same protocol but without *Chlamydia* infectious particles. Cells were fixed using ice cold 100% methanol for one minute. Cells were then blocked with 5% BSA for one hour at room temperature. Focal adhesions and TarP were visualized respectively using a primary monoclonal Talin1 antibody (Novus Biologics, NBP2-50320) and rabbit polyclonal TarP antibody generated against the epitope (661-710 a.a.) (Li International, Denver, CO). Samples were incubated overnight at 4°C. Immunostaining with secondary antibodies was as described above.

### Time-lapse microscopy

For live-cell imaging of FAs fibroblasts were seeded on ibidi µ-slide 8 well chambers with fibronectin coating (ibidi, 80823) at the recommended seeding density and left overnight in a 37 °C, 5% CO_2_ incubator. The following day, cells were infected with CtrL2 with a MOI of 5. At 2 h post-infection, the cells were transfected with either Vinculin-venus (Grashoff *et al*., 2010) a gift from Martin Schwartz, (Addgene, 27300), paxillin-pEGFP (Laukaitis *et al*., 2001) a gift from Rick Horwitz, (Addgene, 15233), or FAK-GFP (Gu *et al*., 1999; Lane *et al*., 2008) a gift from Kenneth Yamada, (Addgene, 50515) using Lipofectamine 3000 transfection reagent (Thermofisher Scientific, L3000008), following the manufacture instructions. After 20 to 22 h, time lapsed images of transfected cells were obtained using a Leica SD6000 AF in TIRF mode, in Washington State University IPN advance image equipment. Images of the GFP-tagged proteins were collected every minute for 90 min. The time lapse images were uploaded to the Focal adhesion Analysis server (Berginski and Gomez, 2013).

### Cell Motility Assay

Mouse embryonic fibroblasts (MEFs) were seeded on ibidi µ-slide live cell imaging chambers (ibidi, 80426). Cells were infected with *Chlamydia trachomatis* serovar L2 at an MOI of 10 by rocking at 4°C for one hour. Infected cells were imaged starting at 20 hours post-infection. A mock-infected control was made following the same protocol but without *Chlamydia* infectious particles. Cells were transfected with N1-mTurquoise2 empty vector control or TarP^829-1006^-mTurquoise2 (LDVBD) using electroporation, seeded into an ibidi µ-slide, and imaged starting at 22 hours post electroporation. Cells were imaged in live cell imaging solution (Thermofisher scientific, A14291DJ) supplemented with 5% fetal bovine serum within a 37°C, 5% CO_2_ controlled environment. Time-lapse DIC images were obtained using a Leica SD6000 AF microscope every ten minutes for ten hours. To minimize the risk of phototoxicity, we restricted image acquisition of the fluorescent mTurquoise2 channel for our transfected cells to the last frame alone. We utilized a similar individual cell tracking data analysis approach as described in (Pijuan *et al*., 2019). Cell motility was tracked using the manual tracking function in ImageJ. Each individual cell was tracked using the position of the nucleus over time. We maximized the time of analysis for each experiment based on the number of cells that remained within a trackable field of view over the imaging span. The coordinate data from the manual tracking function was uploaded to ibidi’s chemotaxis and migration tool. Measurements were taken from spatially calibrated images with a (pixel/µm) scale contained within the meta data. The x/y calibration was set to 0.800001 based on the microscope’s settings contained within the file’s meta data. The statistics function was used to determine the velocity and euclidean (straight-line) distance traveled for each cell.

### *De novo* protein inhibition

Cos7 cells were cultured as previously described. Prior to infection, cells and EB particles were treated with 60 μg/ml of chloramphenicol (Sigma C0378) for 30 min. Cells and EBs were kept in chloramphenicol supplemented DMEM until fixation. Cells were fixed at 8 or 20 h post-infection and were immunostained as described above.

### Western blot

Cos7 cells were plated in 6 well plates with duplicate wells and incubated at 37°C and 5% CO2, until 80% confluency. Cells were infected, as previously described, with CtrL2 for 0 min, 8 or 24 hrs with an MOI of 200. Proteins were harvested using ice cold RIPA buffer (Millipore 20-188) supplemented with phosphatase (Sigma 4906845001) and protease inhibitors (Sigma 5892970001). Cells were scraped and incubated for 30 min on ice. The lysates were centrifuged at 13,000 x g for 20 min at 4°C. Supernatants were diluted in Laemmli buffer (Biorad 161-0747) and kept at −20°C before analysis. Samples were resolved on 10% acrylamide SDS-PAGE. Proteins were transferred to a nitrocellulose membrane (Bio-rad 1620115). Immunoblotting was performed by blocking membranes with 5% BSA in TBS-T overnight at 4°C and incubation using antibodies against TarP (a generous gift from Dr. Raphael Valdivia, Duke University) and β-tubulin HRP conjugated (Abcam ab21058). The secondary antibody used was anti-mouse HRP conjugated (DAKO P0161). Immobilin chemiluminescence kit (Millipore, WBKLS0500) was used to develop the blot.

### Cloning and transfection of TarP contructs

A summary of the primers used in this study is provided in Table S1. Initially TarP Full-length, TarP LDVBD, TarP LD, and TarP VBD were PCR amplified from CtrL2 genomic DNA using the primers combination 1-2, 5-2, 5-6 and 4-2, respectively. A *Bam*H1 (reverse primer) and *Kpn*I (forward primer) restriction sites were used for fusion with the N1-mTurquoise2 plasmid. The TarP *Δ*PRD was obtained using the 7-8 primer pair for PCR amplification from TarP Full-length-mTurquoise2 fusion plasmid. The primers were created to amplify the whole TarP Full-length-mTurquoise2 except the nucleotides that constitute the proline rich domain (PRD) 625-650. The resulting PCR product was recombined using in-Fusion HD cloning plus CE (Clontech, 638916) to create a functional circular plasmid. TarP *Δ*LDVBD was PCR amplified from the TarP *Δ*PRD-mTurquoise2 plasmid using the primers pair 1-3. The same restriction enzymes were used to clone these fragments into N1-mTurquoise2. Transformations using restriction enzymes recombination were made into chemically competent Top10 (invitrogen) *E. coli*, and vectors sequence was verified using sequencing (Eurofins) The construct pFH-TarP *Δ*PRD and pFH-TarP *Δ*LDVBD used for super-resolution experiments was PCR amplified from TarP *Δ*PRD-mTurquoise2 using primers combination 9-10 and 9-17, respectively. To use homology cloning the vector backbone 1436 pcDNA3-Flag-HA, kindly provided by William Sellers (Addgene 10792), was linearized by PCR using the primers pair 11-12 and 11-18, creating homology overhang regions to the TarP *Δ*PRD and TarP *Δ*LDVBD, respectively. TarP LDVBD was amplified from CtrL2 genomic DNA using the primer pairs 13-14. To use homology cloning the vector backbone 1436 pcDNA3-Flag-HA was linearized by PCR using the primer pair 15-16. Fragments and vector backbone were recombined using in-Fusion HD cloning plus CE (Clontech, 638916) to create a functional circular plasmid. Transformations using homology recombination were made into chemically competent Stellar (Clontech) *E. coli*, and vectors sequence was verified using sequencing (Eurofins). The pcDNA3-Flag-Apex-Nes was a gift from Alice Ting (Addgene, 49386). The N1-mTurquoise2 (Addgene, 54843). Transfections were done as described above. For iPALM experiments 1µg of DNA and 2µl of sheared salmon sperm DNA were mixed together in 15µl of Opti-MEM (Thermofisher scientific, 31985062), and kept on ice for 15 min. 1×10^6^ Cos7 cells were resuspended in 200µl of cold Opti-MEM, mixed with the DNA solution and kept on ice for 30 seconds. Cells and DNA suspension were transferred to a 4mm gap cuvette (BioRad, 1652088) and electroporated using BioRad Gene Pulser XCell using the following settings: 190V; 950uF; infinity. After electroporation 1.5ml of warm growth media was added. 400µl of cell solution was added to a 6 well plate well containing 1.5ml of warm growth media and the gold fiducial coverslip. Cells were incubated 37°C, 4% CO_2_ for 4 h to adhere to the gold fiducial coverslip. Cells were washed to remove dead cells debris and further incubated for 20 h.

### iPALM imaging and analysis

The principle of instrumentation for iPALM imaging and analysis were performed as previously described (Shtengel *et al*., 2009; Kanchanawong *et al*., 2010) with the following modifications. After 24°C of transfection cells plated in gold fiducial coverslip were fixed with 0.8% PFA and 0.1% glutaraldehyde (Sigma G7526-10ML) solution (in PBS) for 10 min. After fixation cells were washed 3 times with PBS and quenched using 1% NaBH4 (Sigma, 452882-25G) solution (in PBS) for 7 min. Cells were then washed again 3 times with PBS. After washing cells were immunostained (when necessary) and/or processed for iPALM imaging as previously described (Kanchanawong *et al*., 2010). The vertical coordinates relative to the golden fiducial markers are indicating by a color scale from red (0 nm) to purple (250 nm).

### Trypsin assay

Cells were plated in 24 well plates and incubated at 37°C and 5% CO2, until 80-90% confluency. Afterwards, cells were infected with CtrL2 with a multiplicity of infection of 5 for 20 h. Cells were then treated with 0.01% trypsin diluted in serum-free DMEM media at 37°C, for 0, 10, 20, 30, or 35 min. Cells were fixed with 4% PFA, carefully washed with PBS and stained with DAPI to count the number of remaining cells as well as to visualize *Chlamydia* inclusions. Images were taken using Nikon eclipse TE2000-U. Detachment assays were performed as above, and transfection procedures were as described.

## Acknowledgments

We would like to thank Drs. Martin Schwartz, Rick Horwitz, Kenneth Yamada, William Sellers, Alice Ting, and Michael Davidson for various constructs obtained through Addgene; Dr. Raphael Valdivia (Duke University Medical Center) for the generous gift of the TarP antibody; Dr. Lisa Rucks (University of Nebraska Medical Center) for the *C. trachomatis* serovar D strain; the services provided by the WSU Integrated Physiology and Neuroscience imaging facilities; and Satya Khuon (AIC Janelia) for technical help and advice. This project was supported by grants from the National Institute for Food and Agriculture (#1010265), National Institutes of Health (AI065545), and start-up funds from the WSU College of Veterinary Medicine to RAC. ATP and ATN were recipients of the Fundação para a Ciência e Tecnologia, SFRH/BD/76741/2011 and SFRH/BD/86670/2012, respectively. TRT is a Medical Research Council studentship awardee. KNM was supported by the NIH Protein Biotechnology Training Grant T32 GM008336 awarded to WSU. AJB was supported by the Infectious Diseases and Microbial Immunology Training Grant T32 AI007025 awarded to WSU. iPALM data used in this publication was produced in collaboration with the Advanced Imaging Center, a facility jointly supported by the Gordon and Betty Moore Foundation and Howard Hughes Medical Institute at the Janelia Research Campus.

## Author Contributions

Conceptualization, RAC; Methodology, ATP, JA, and RAC; Software, JA; Formal Analysis, ATP; Investigation, ATP, ATN, KNM, JA, AJB, and TRT; Writing – Original Draft, ATP, and RAC; Writing – Review & Editing, RAC, KNM and AJB; Funding Acquisition RAC; Resources, JA, TLC and RAC; Supervision, TLC and RAC.

---

**Supplemental Table 1.**
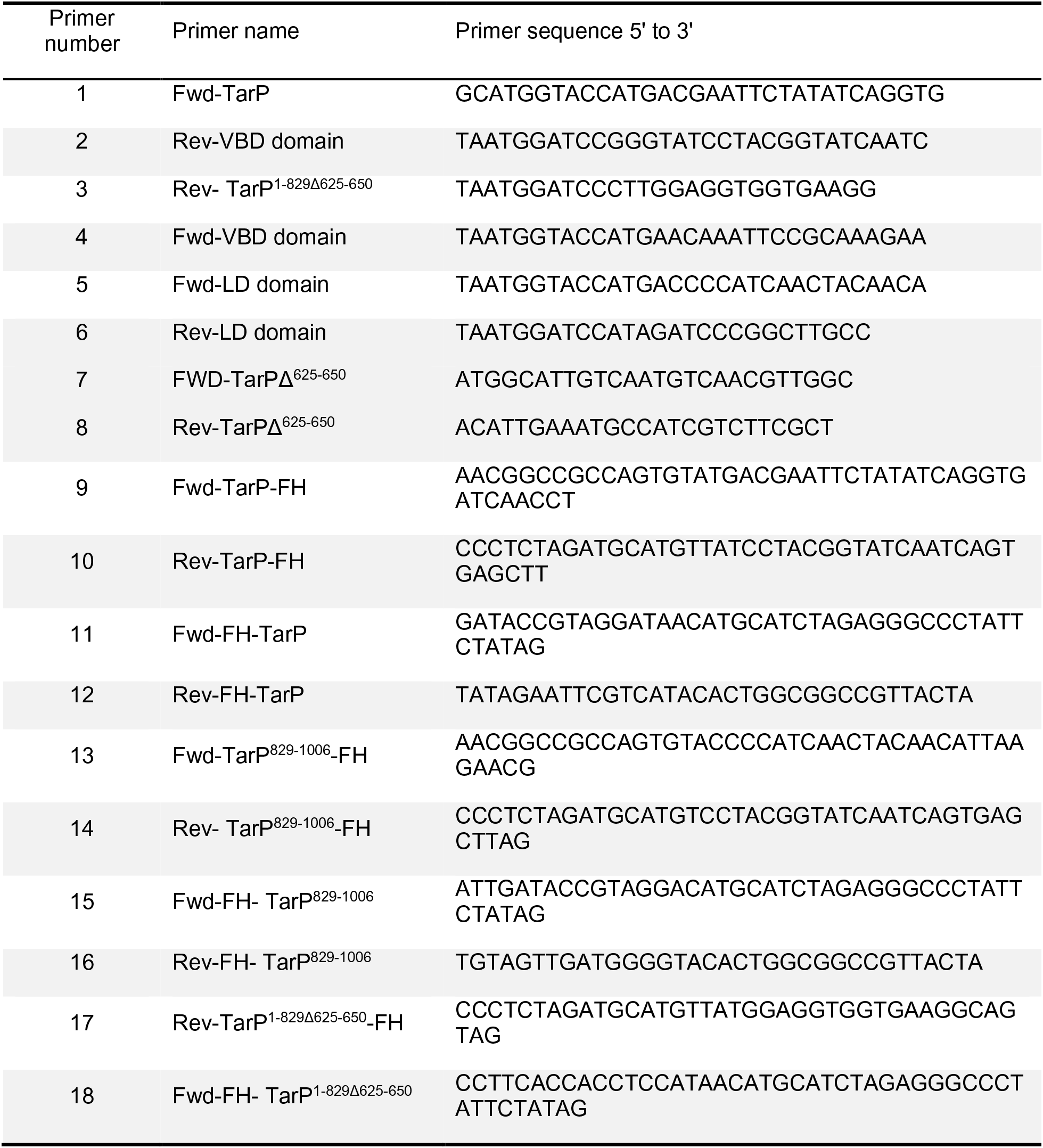
Primers used in this study

**Figure S1.**
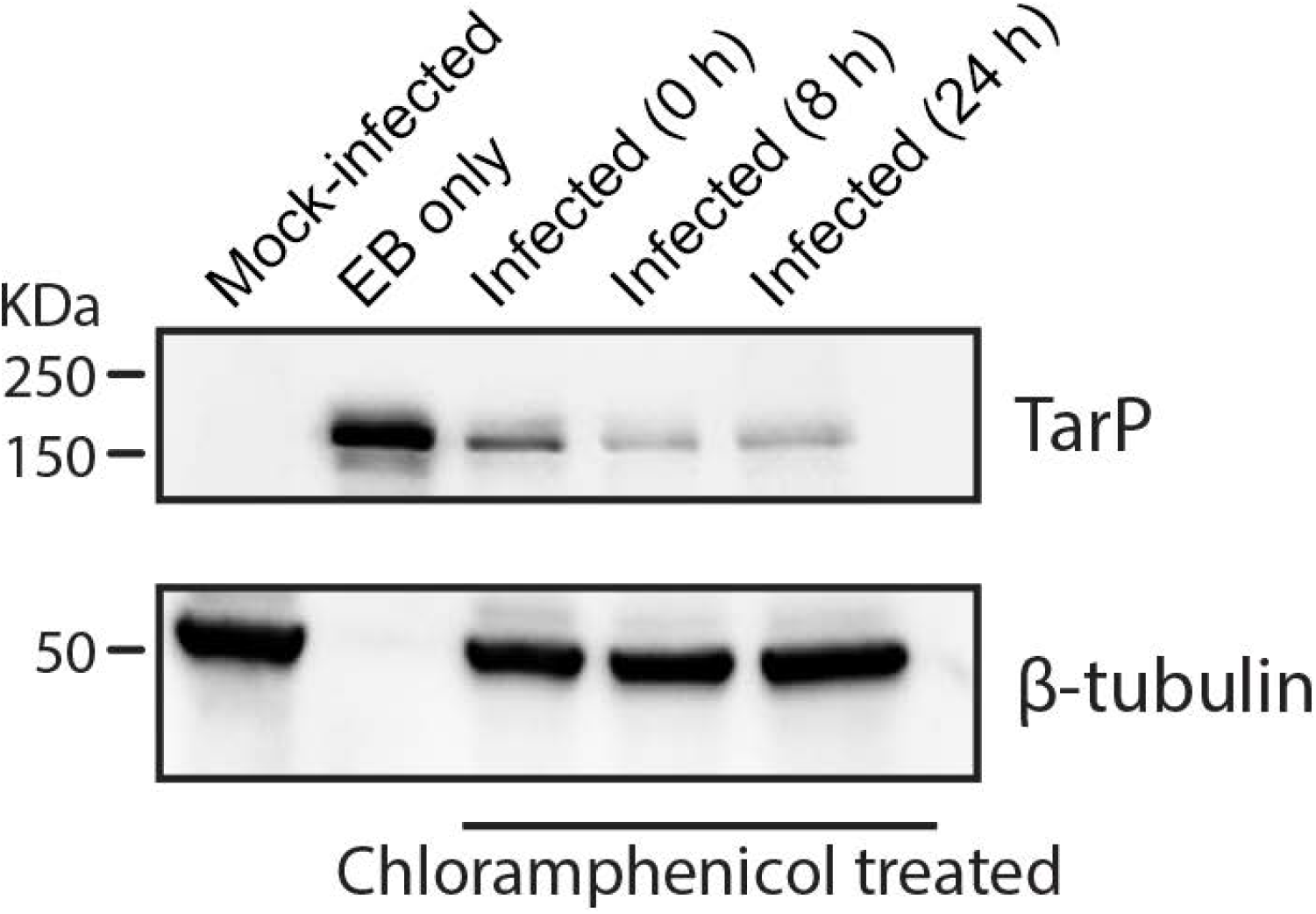
TarP translocated by *C. trachomatis* is stable. (A) Cos7 cells were either mock-infected or *Chlamydia*-infected for the indicated times, and maintained throughout in the presence of the prokaryotic protein synthesis inhibitor chloramphenicol. Whole cell lysates were harvested to monitor by Western blot the presence and stability of translocated TarP proteins. A rabbit polyclonal anti-TarP antibody was used. Protein concentrations were determined and adjusted to ensure equal loading. β-tubulin was used as the loading control. Mock-infected and EB-only samples were added to demonstrate specificity of the anti-TarP antibody used.

**Figure S2.**
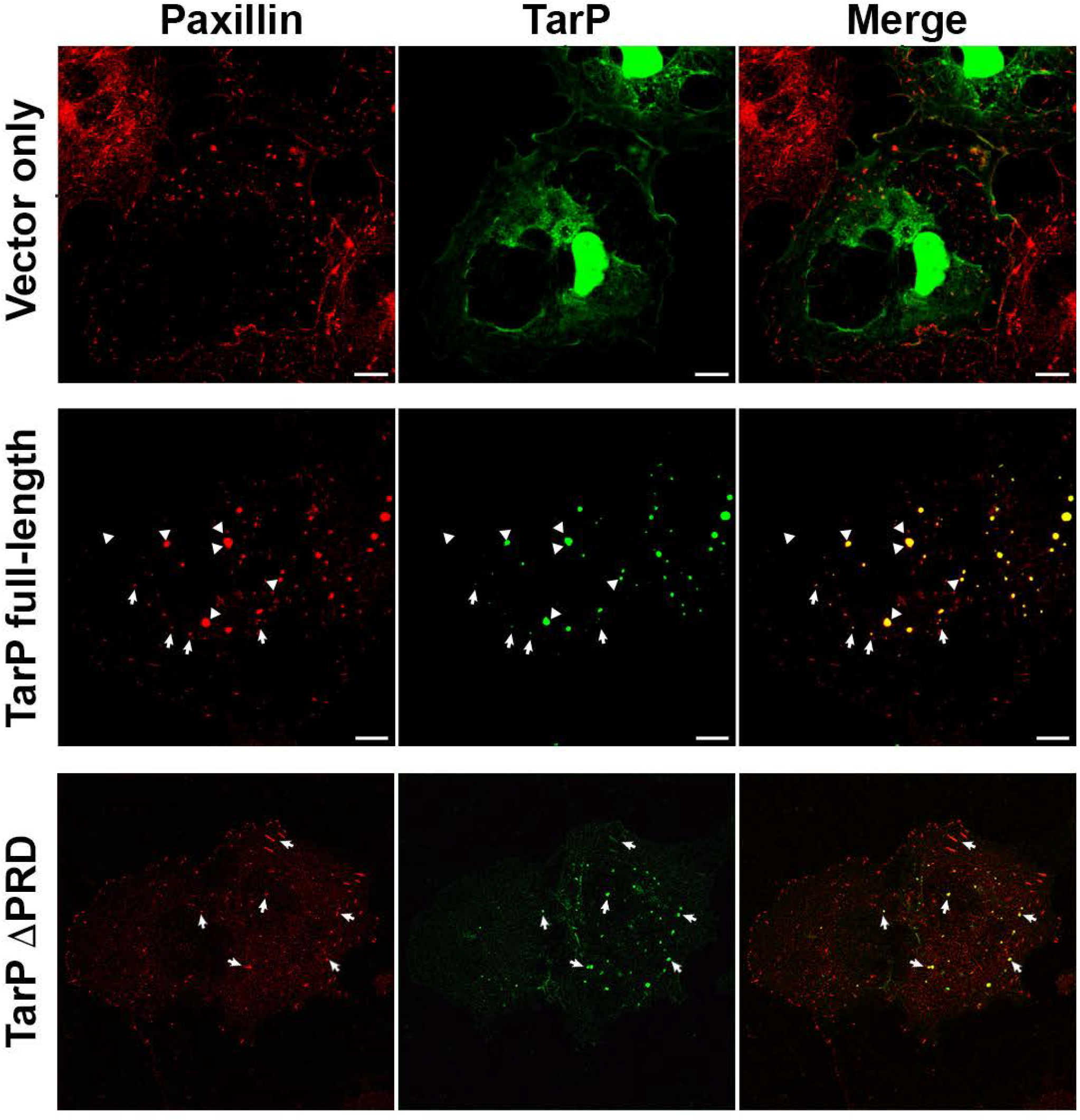
The PRD domain of TarP is not required for focal adhesion localization upon ectopic expression. Full-length TarP and TarP ΔPRD, both fused to mTurquoise2 were expressed ectopically in wild type MEFs for 19 h. The cells were processed for immunostaining for paxillin to visualize focal adhesions. Both full-length TarP and TarP ΔPRD colocalized with paxillin-positive structures, with the forming forming larger protein aggregates (arrowheads). Smaller TarP-positive punctae are indicated by white arrows.

**Figure S3.**
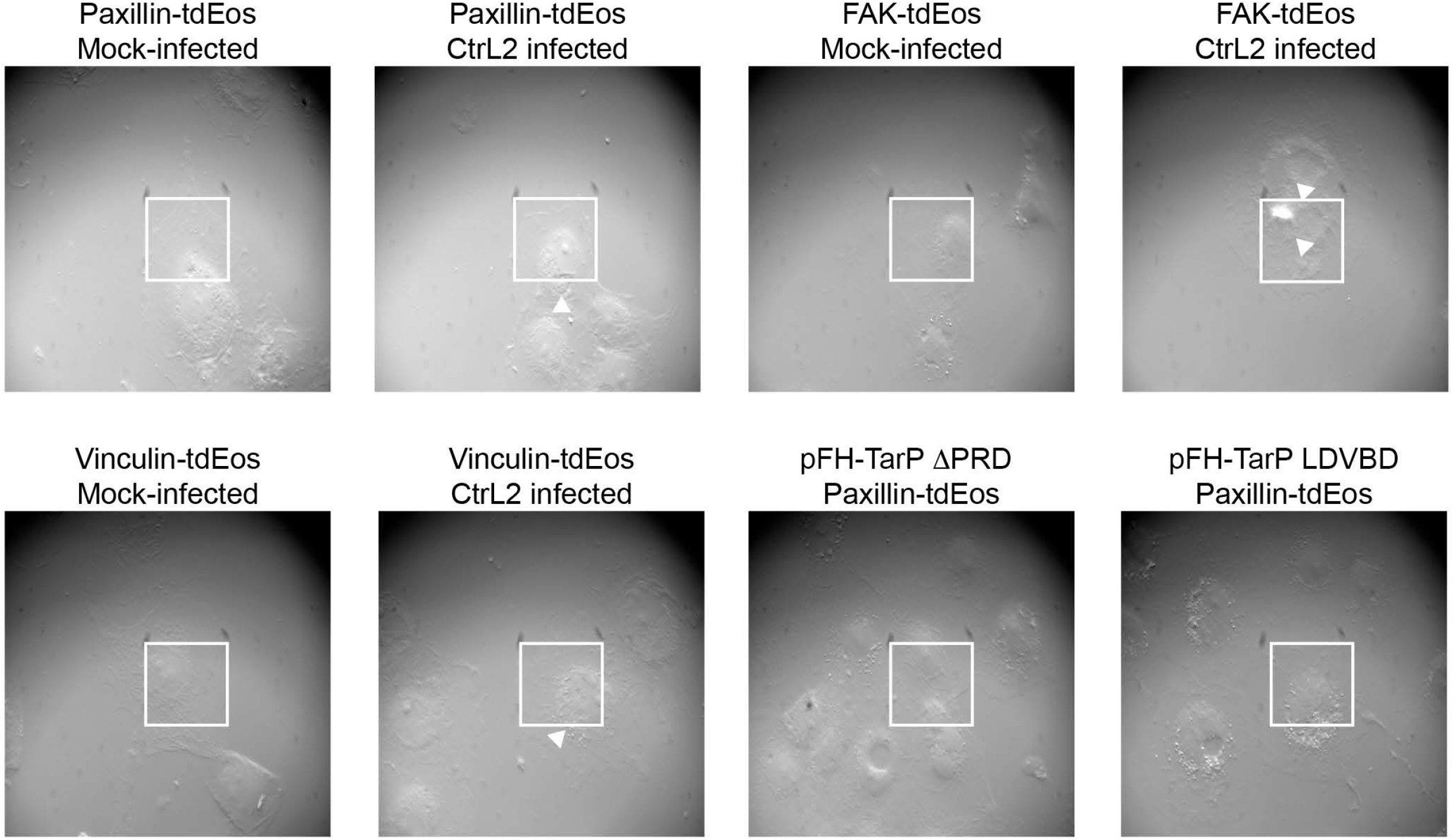
Infection disrupts the stratified organization of focal adhesions. Cos7 cells were either mock-infected or infected with *C. trachomatis* serovar L2 for 20 h. DIC images of cells analyzed by iPALM were acquired to demonstrate the infection state. Regions of interests are bounded by white lines, and inclusions indicated by arrowheads.

**Figure S4.**
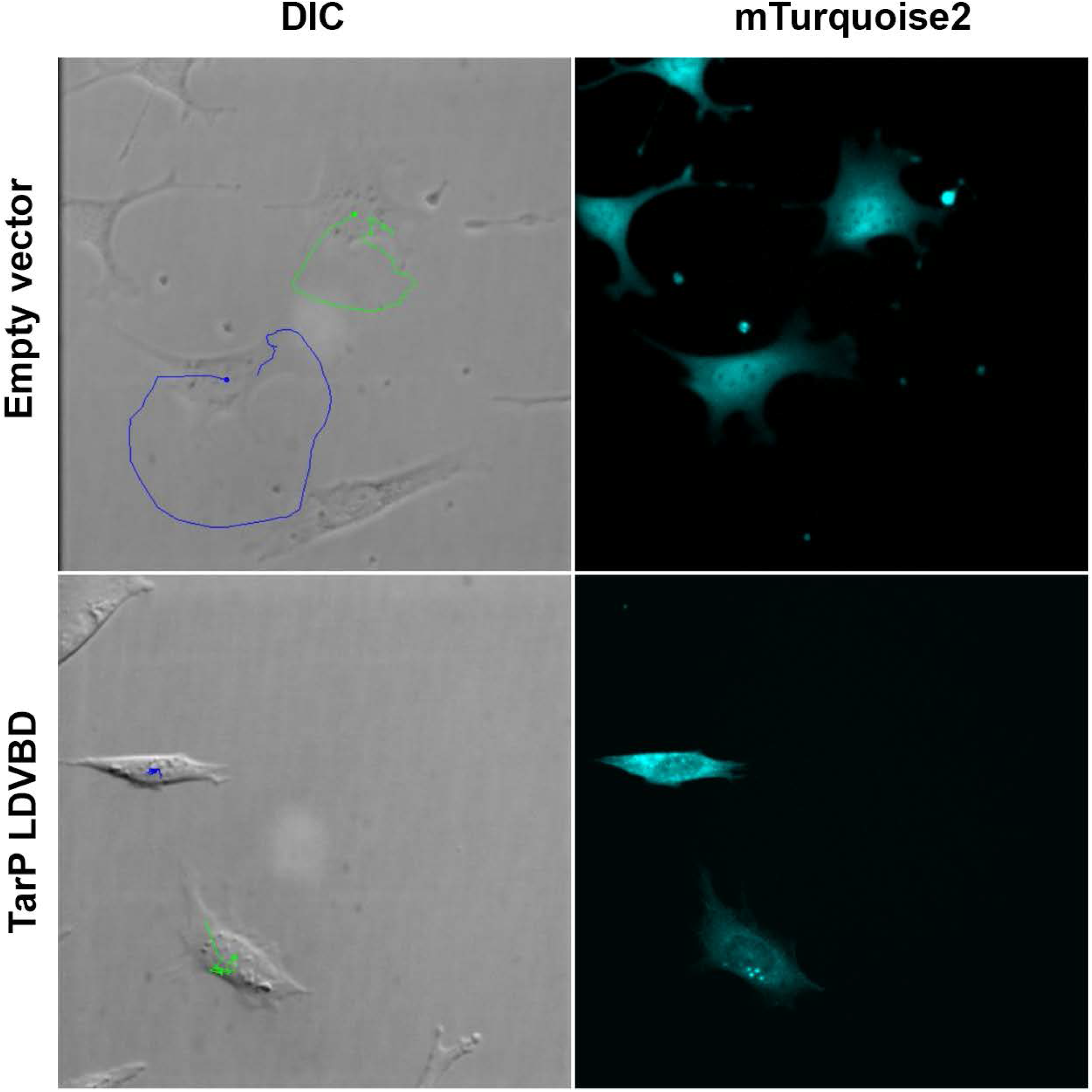
LDVBD-expressing cells exhibited restricted motility relative to the empty vector-transfected control cells. Acquisition of the fluorescent channel was limited to the final frame of the motility assay to minimize phototoxicity. Images show the final DIC and fluorescent channel captured following 10 hours of time-lapse imaging. The DIC channel includes the dot and line overlay generated via ImageJ’s manual tracking function to indicate the cell’s movement over time.

**Movie S1. Infection inhibits cell motility.** A representative video assembled from a 17-h time-lapse imaging of mock-infected MEFs shows motility of individual cells with track outlines included.

**Movie S2. Infection inhibits cell motility.** A representative video assembled from a 17-h time-lapse imaging of *C. trachomatis*-infected MEFs shows motility of individual cells with track outlines included.

**Movie S3. LDVBD is sufficient to inhibit cell motility.** MEFs transfected with the vector alone was monitored for 17 h. A representative video assembled from a series of time-lapse images shows the degree of cell motility in the vector-only control group.

**Movie S4. LDVBD is sufficient to inhibit cell motility.** MEFs transfected with the LDVBD-mTurquoise2 was monitored for 17 h. A representative video assembled from a series of time-lapse images shows inhibition of cell motility of LDVBD-expressing cells.

**Movie S5. *Chlamydia*-infected cells are resistant to detachment by mild trypsinization.** Mock-infected HeLa cells growing on glass coverslips were treated by 0.025% Trypsin + EDTA, and imaged at 60-s intervals for 30 min. Note that the cells start to round up by seven min of incubation in trypsin.

**Movie S6. *Chlamydia*-infected cells are resistant to detachment by mild trypsinization.** *C. trachomatis* serovar L2-infected HeLa cells growing on glass coverslips were treated by 0.025% Trypsin + EDTA at 24 h post-infection, and imaged at 60-s intervals for 30 min. In contrast to mock-infected cells shown in Movie S5, the infected cells remained attached and spread out after 30 min of mild trypsinization, indicating a possible enhancement of adhesion of infected cells to the substrate.

